# The MYCN oncoprotein and helicases DDX17 and DDX5 have opposite effects on the production of chimeric transcripts in neuroblastoma cells

**DOI:** 10.1101/2024.02.05.578895

**Authors:** Valentine Clerc, Jessica Valat, Xavier Grand, Nicolas Fontrodona, Matéo Bazire, Nicolas Rama, Didier Auboeuf, Benjamin Gibert, Franck Mortreux, Cyril F. Bourgeois

**Author notes:** **Corresponding author**: Cyril F. Bourgeois.

## Abstract

DEAD box helicases DDX17 and DDX5 control the termination of transcription and the associated cleavage of the 3’ end of transcripts. Here we show that the transcriptional readthrough induced by their depletion in neuroblastoma cells also results in increased production of chimeric transcripts from tandemly oriented genes. Analysis of neuroblastoma tumours in which chimeric transcripts are abundant revealed that low expression of the DDX17 and DDX5 genes is associated with poor overall patient survival. Low DDX17 expression is also significantly associated with high-risk tumours and is inversely correlated with MYCN oncogene amplification, suggesting a link between these two factors. We demonstrate that changes in MYCN expression do not affect the expression of either helicase, but alter transcription termination leading to the production of chimeric transcripts. We provide evidence that MYCN acts on termination through its direct binding to the 3’ region of genes and that it interacts with DDX17, suggesting that it may inhibit the activity of the helicase. Collectively, our work reveals a novel function of MYCN in transcription termination and suggests that the deregulation of MYCN and DDX17/DDX5 expression in neuroblastoma may lead to the expression of non-canonical and potentially harmful RNA molecules.

## Introduction

During transcription by RNA polymerase II (RNAPII) in eukaryotes, the nascent RNA undergoes different processing steps, including 5’ capping, splicing, 3’ end cleavage and addition of the poly-A tail. These steps are mostly co-transcriptional and are impacted not only by RNA binding proteins, but also by transcription and chromatin-associated factors (Neugebauer, 2019, Tellier et al., 2020). The tight connection that exists between transcription termination and 3’ end RNA processing is archetypal of this coordination of events. The slow down and release of RNAPII after the transcription of the polyadenylation site (PAS) and the cleavage of the nascent transcript are mutually dependent on each other, in a complex interplay (Boreikaite and Passmore, 2023, Porrua and Libri, 2015, Proudfoot, 2016, Rodriguez-Molina et al., 2023). The regulation of this process involves a number of factors that act on chromatin, at the level of the RNAPII complex or on the RNA molecule, or which can have an effect at several levels.

An alteration in the expression or activity of factors involved in transcription termination and/or RNA cleavage classically results in transcriptional readthrough and in the extension of the 3’ extremity of the transcript. Readthrough transcription has been observed in cells exposed to various stresses or viral infection, as well as in cancer (Cardiello et al., 2018, Grosso et al., 2015, Rutkowski et al., 2015, Heinz et al., 2018, Rosa-Mercado et al., 2021, Vilborg et al., 2015, Vilborg et al., 2017, Hennig et al., 2018, Bauer et al., 2018), as recently reviewed (Rosa-Mercado and Steitz, 2022, Morgan et al., 2022). The fate of these extended transcripts is uncertain, but it has been shown that stress-induced DoGs (downstream of gene containing transcripts) lack a defined endpoint and are non-coding RNAs which remain associated to the chromatin in the nucleus (Vilborg et al., 2015, Vilborg et al., 2017).

In some cases, RNAPII that fails to terminate past the PAS can invade the downstream gene on the same DNA strand, generating *cis*-spliced transcripts containing exons from both genes. These RNA molecules have been called transcription-induced chimeras (Akiva et al., 2006, Parra et al., 2006), tandem RNA chimeras (Greger et al., 2014), products of conjoined genes (Prakash et al., 2010, Kim et al., 2012), readthrough fusions (Varley et al., 2014) or *cis*-splicing products of adjacent genes (Qin et al., 2016, Qin et al., 2015). Below we will simply define them as chimeric transcripts, keeping in mind that in this study we are not considering RNA molecules resulting from chromosomal rearrangements. The formation of chimeric transcripts can be associated to splicing defects in the adjacent invaded gene, highlighting the necessary coordination of events that ensures the correct and autonomous expression of neighbouring genes (Alpert et al., 2020, Hadar et al., 2022). Thanks to the increasing number and deeper analysis of RNA-seq experiments, chimeric transcripts have been identified in many types of cancers and associated with oncogenesis (Dorney et al., 2023, Sun and Li, 2022). However, they are also found in normal tissues (Babiceanu et al., 2016) and their production is considered as a possible mechanism to increase protein diversity (Parra et al., 2006). Indeed, chimeric transcripts can retain a functional open reading frame (ORF) allowing the production of chimeric or fusion proteins (Varley et al., 2014, Yun et al., 2014, Han et al., 2017, Egashira et al., 2019).

Several factors have been shown to affect the formation of chimeric transcripts in human cells, including splicing and/or polyadenylation factors (Chwalenia et al., 2019). Chemical splicing inhibition represses the formation of chimeric transcripts in neuroblastoma, a cancer in which most RNA fusions are intrachromosomal, likely resulting from transcriptional readthrough and *cis*-splicing (Shi et al., 2021). Chromatin-associated factors are also involved in chimeric RNA production, such as the histone methyltransferase SETD2 (Grosso et al., 2015) or the CCCTC-binding factor CTCF (Qin et al., 2016, Qin et al., 2015), which plays important roles in spatial genome organization and gene expression regulation via chromatin looping or insulation (Braccioli and de Wit, 2019). Recent reports further supported a function of CTCF in controlling PAS choice and transcription termination (Nanavaty et al., 2020, Terrone et al., 2022).

DEAD box ATP-dependent RNA helicase DDX17 and its paralog DDX5 have a variety of functions in gene expression, especially in transcription and RNA processing (Giraud et al., 2018, Xing et al., 2019). Thanks to their helicase activity, they act by modulating local RNA secondary structures or DNA/RNA structures such as R-loops and G-quartets, impacting promoter and exon choice (Camats et al., 2008, Kar et al., 2011, Lai et al., 2019, Mersaoui et al., 2019, Wu et al., 2019). Recently, we and others have shown that DDX17 and DDX5 control the 3’ end processing of RNAPII transcripts, and that their knockdown leads to a decreased termination downstream of the expected PAS (Katahira et al., 2019, Lai et al., 2019, Mersaoui et al., 2019, Terrone et al., 2022). We showed that this function is linked to the binding of CTCF near the PAS and to the specific 3D organization of their target genes (Terrone et al., 2022). We hypothesized that the helicases could control the dynamics of transcription and RNA processing across DNA and/or RNA structured regions, in line with other reports that described an impact on R-loops or 3’UTR RNA structure (Katahira et al., 2019, Lai et al., 2019, Mersaoui et al., 2019). We show hereafter that the transcription readthrough induced by *DDX17* and *DDX5* silencing in neuroblastoma cells results in the production of several hundreds of chimeric transcripts, a fraction of which is also found in neuroblastoma tumours. Interestingly, we found that high *DDX17* and *DDX5* expression correlates with good survival of neuroblastoma patients and that it is inversely correlated to the amplification of *MYCN*, a driving oncogene in this cancer (Otte et al., 2020). Like the closely related MYC protein, MYCN is a transcription factor which binds to most active promoters and regulates several aspect of RNAPII activity (Baluapuri et al., 2020). Here, we present evidence supporting a direct role of MYCN in controlling transcription termination through its binding near the PAS of DDX17/DDX5-targeted genes. This new function of MYCN suggests that the combined amplification of this oncogene and suboptimal expression of *DDX17* and *DDX5* could be determinant to explain the increased expression of chimeric transcripts in neuroblastomas.

## Results

### DDX17 and DDX5 depletion enhances the expression of chimeric transcripts

We recently described a genome-wide effect of DDX17/DDX5 depletion on transcription termination in neuroblastoma cells SH-SY5Y cells (Terrone et al., 2022). In our RNA-seq results, we observed numerous examples of transcriptional readthrough induced by DDX17/DDX5 depletion that also displayed splicing junction reads across adjacent genes. Typically, splicing skipped the terminal exon of the altered gene and linked its penultimate exon to the second exon of the following gene, generating a chimeric mRNA molecule (Fig. 1A, Supp. Fig. 1A). However, we also frequently observed complex alternative splicing patterns across exons from both genes (Supp. Fig. 1B). Interestingly, in some cases chimeric junctions linked more than 2 adjacent genes, suggesting that DDX17/DDX5 depletion could induce the formation of multimeric mRNA molecules (*NTRK1-PEAR1-LRRC71* and *NDUFA13-YJEFN2-CILP2*, Supp. Fig. 1B).

**Figure 1.**
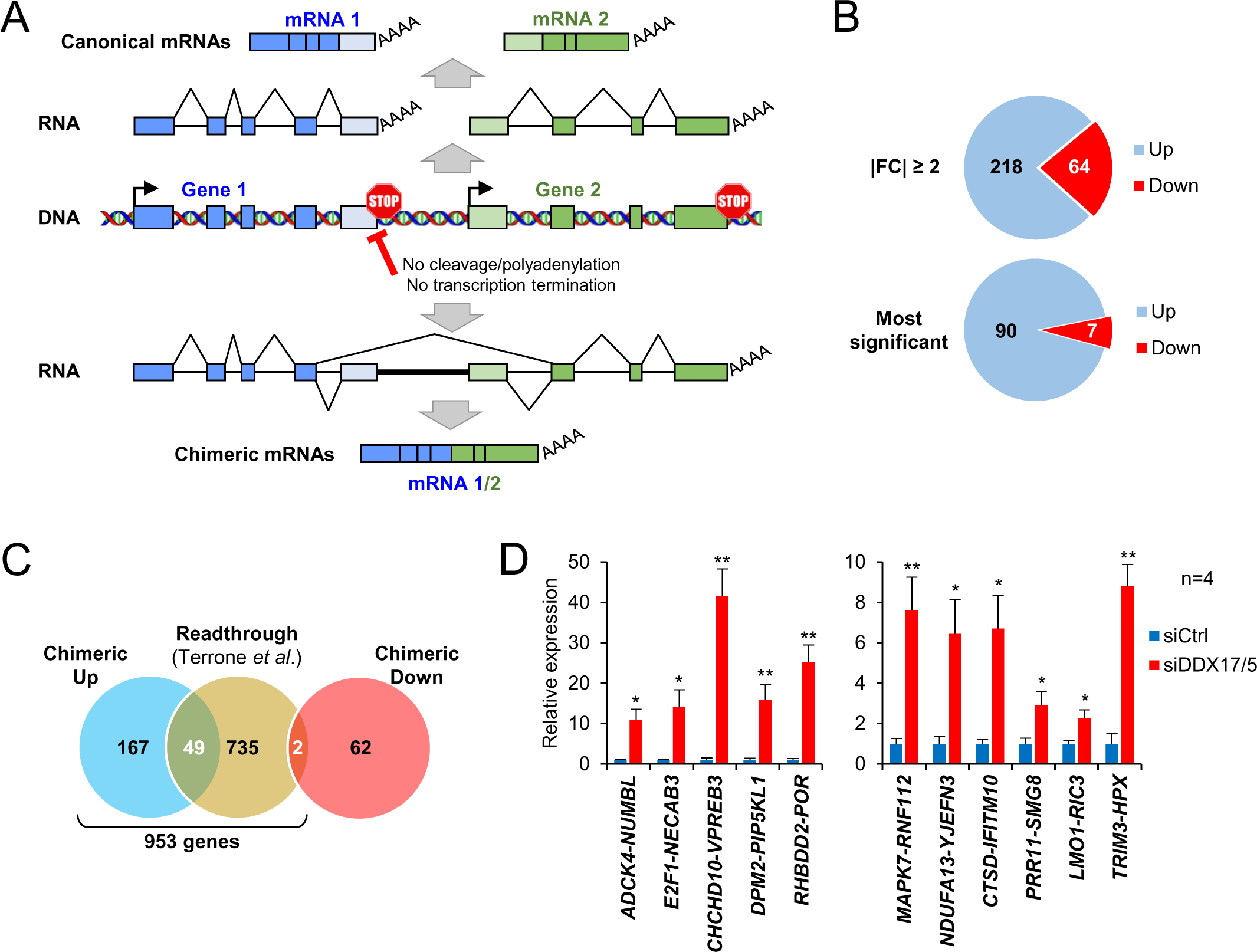
DDX17 and DDX5 depletion enhances the expression of chimeric transcripts. **A.** Schematic representation of the production of readthrough induced chimeric transcripts. **B.** Diagram showing the number of up and downregulated chimeric trancripts displaying at least a fold change of 2 and the most significant transcripts (*P*-val<0.1). **C.** Venn diagram showing the overlap between genes producing chimeric transcripts and the genes previously identified as displaying transcriptional readthrough events. The list of 953 genes corresponds to the genes on which we carried out global analyses (Supp. Table 1). Note that among the 218 upregulated chimeric transcripts from panel B, there are 2 cases in which one exon from the gene of origin is spliced with exons from different target genes, generating 2 different chimeras. As these events can be counted only once in panel B, this explains the total of 216 upregulated chimeras. **D.** Quantification of chimeric transcripts. Expression of chimeric transcripts was monitored by RT-qPCR using primers spanning the chimeric junction and then normalized on the expression of the 5’ parental gene (see also Supp. Fig. 2B). Two-tailed paired t-test (* *P*-val<0.05 ; ** *P*-val<0.01).

We next used the previously described Arriba algorithm (Uhrig et al., 2021) to determine more precisely the number of chimeric transcripts whose expression is modified upon DDX17/DDX5 depletion. We identified 282 chimeric mRNAs, 218 of which (77%) were upregulated and 64 downregulated (Fig. 1B, top panel, Supp. Table 1). Applying a more stringent selection threshold confirmed that the depletion of both helicases indeed mostly increased the expression of chimeric transcripts (Fig. 1B, bottom panel). We found that a large proportion of the genes displaying an induced chimeric transcript had been previously identified as genes with readthrough induced by siDDX17/DDX5 (Fig. 1C) (Terrone et al., 2022). From these analyses, we defined a list of 953 genes whose termination is inhibited upon depletion of DDX17 and DDX5 (Supp. Table 1).

We next sought to experimentally validate the expression of a selection of chimeric transcripts. We first carried out RT-PCR experiments using a primer overlapping the RNA fusion between both genes, to specifically detect the chimeric transcript, that we systematically compared to the canonical form of the gene. We observed an increased amount of all chimeric transcripts tested upon DDX17/DDX5 depletion, which was not consistent with the expression of the corresponding canonical mRNA (Supp. Fig. 2A). We confirmed this result by RT-qPCR on a panel of chimeric transcripts, demonstrating that the normalized expression of all molecules was increased between 2 and 40-fold in absence of DDX17/DDX5, compared to the canonical transcript (Fig. 1D and Supp. Fig. 2B). Note that validated *CTSD-IFITM10*, *NDUFA13-YJEFN2* and *TRIM3-HPX* chimeric RNAs were not predicted as significantly induced by Arriba but were identified while browsing through our RNA-seq data (Supp. Fig. 1A-B). This suggested that the actual number of chimeric RNAs resulting from DDX17/DDX5 depletion is likely underestimated.

### Chimeric transcripts can produce chimeric proteins

Some chimeric transcripts maintain a coding sequence that matches the reading frame of their respective canonical transcripts, potentially allowing them to escape degradation by the nonsense-mediated mRNA decay pathway. For example, the *CTSD-IFITM10* chimer lacks the last exon of *CTSD* and the first exon of *IFITM10*, but it is cleaved and polyadenylated at the canonical PAS of the *IFITM10* gene. This chimeric mRNA is predicted to produce a chimeric protein of 557 amino acids (60 kDa), which lacks 55 and 28 amino acids from the C-terminal part of CTSD and N-terminal part of IFITM10 proteins, respectively (Fig. 2A), and which was identified in breast cancer cells (Varley et al., 2014).

**Figure 2.**
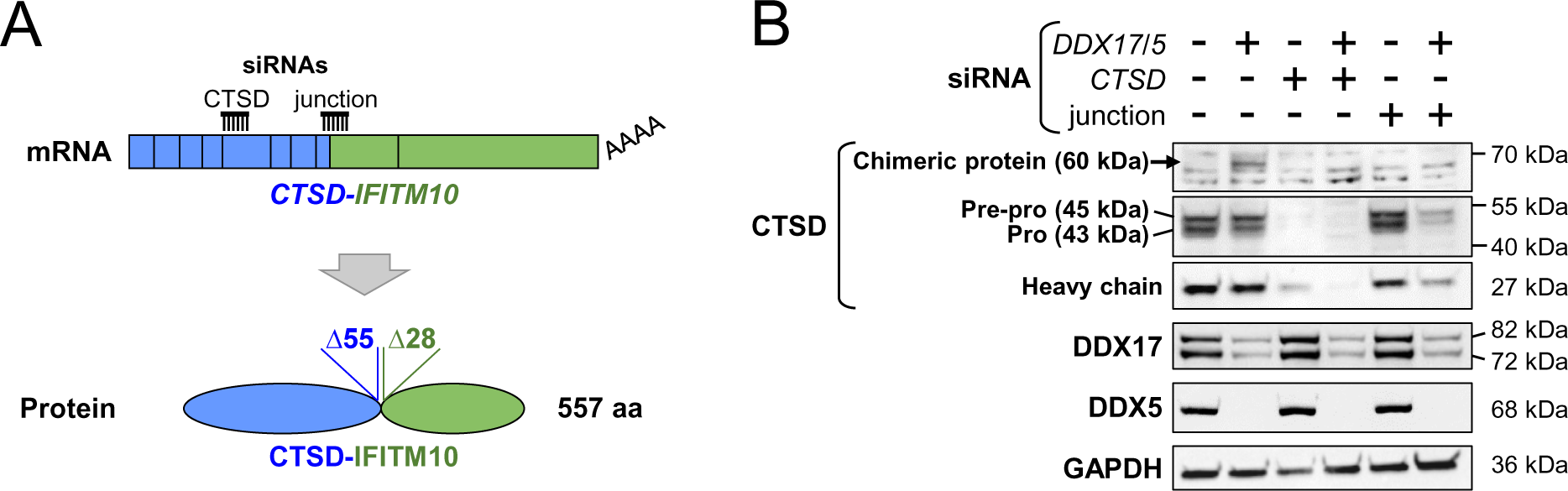
Chimeric transcripts can produce chimeric proteins. **A.** Schematic representation of *CTSD-IFITM10* chimeric transcripts and putative chimeric protein. The sites targeted by the siRNAs used to validate the specificity of the band corresponding to CTSD-IFITM10 protein are indicated. **B.** Western-blot showing the expression of the chimeric CTSD-IFITM10 protein and the canonical protein CTSD, as well as the validation of *DDX5* and *DDX17* silencing.

We carried out a western-blot analysis to detect this chimeric protein in neuroblastoma cells, using an antibody against CTSD. The *CTSD* gene normally encodes a unique precursor that is processed into two peptidic chains that compose Cathepsin D. As expected, we identified the two subunits between 25 and 45 kDa (Fig. 2B), but we also detected a discrete 60 kDa band that appeared only upon depletion of DDX17 and DDX5. All bands disappeared when cells were treated with a siRNA targeting the body of the *CTSD* transcript. Furthermore, treating cells with another siRNA specifically targeting the *CTSD-IFITM10* junction also led to a loss of the 60 kDa band. Note that this treatment also reduced the level of CTSD isoforms, likely because of a non-specific effect of the junction siRNA on canonical mRNAs.

This result demonstrated that the siDDX17/DDX5-induced chimeric mRNAs can produce chimeric proteins, which could have a strong impact on the functions of their respective canonical proteins.

### The expression of the *DDX17* gene is reduced in high-risk neuroblastomas

Recently, neuroblastoma tumours were shown to express large amounts of chimeric transcripts (Shi et al., 2021), which prompted us to look whether DDX17/DDX5-regulated chimeras can be found in these tumours. Of the 97 main chimeric transcripts from our analysis, 25 were also identified in neuroblastoma tumours (Fig. 3A, top panel). All these mRNAs belong to the pool of chimeric transcripts upregulated upon DDX17/DDX5 depletion, while none of the downregulated transcripts was identified (the top 10 transcripts are shown in Fig. 3B, all transcripts are shown in Supp. Table 1). Of note, only 1 of these 25 transcripts (*SLC29A1-HSP90AB1*) was also detected in a control cohort of 161 healthy adrenal gland samples, while 4 of them were detected in at least 10% of neuroblastomas (Shi et al., 2021) (Fig. 3A, bottom panel and Fig. 3B).

**Figure 3.**
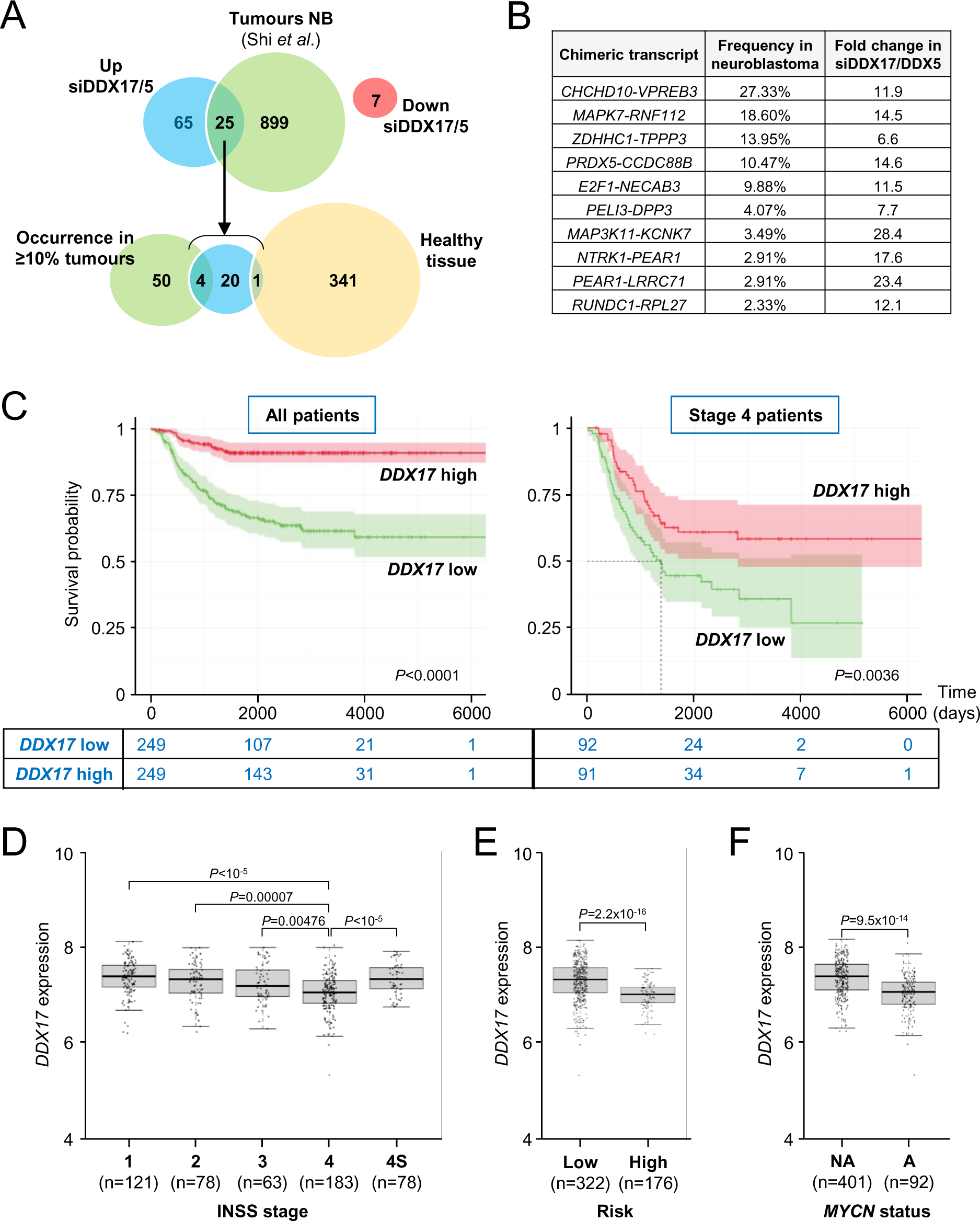
The expression of the *DDX17* gene is reduced in high-risk neuroblastomas. **A.** Venn diagrams showing the number of chimeric transcripts (defined as in Figure 1A) that are both up or down-regulated after DDX17/DDX5 depletion and identified in neuroblastoma tumors. **B.** Top 10 of transcripts identified both in neuroblastoma tumours and upregulated upon DDX17/DDX5 depletion. **C.** Kaplan-Meier curves showing the overall survival of neuroblastoma patients, separated in 2 groups of high and low *DDX17* expression (relative to the median value of the group). Left diagram: all patients (n=498). Right diagram: stage 4 patients (n=183). The number of patients in each group and for each time point is indicated below. **D.** Box plot of *DDX17* expression relative to tumor stage (INSS classification). ANOVA corrected for multiple comparisons with Tukey’s tests. Only the comparison between Stage 4 and other stages is shown, the other comparisons were not significant. **E.** Box plot of *DDX17* expression relative to low/high risk classification. Two-tailed *t*-test. **F.** Box plot of *DDX17* expression relative to amplified (A) or non-amplified (NA) *MYCN* status. Two-tailed *t*-test.

Our earlier work showed that DDX17 is involved in the early phases of retinoic acid-induced differentiation of neuroblastoma cells (Lambert et al., 2018), and recent findings have associated a high expression DDX5 with an unfavourable outcome in neuroblastoma patients (Zhao et al., 2020). To explore further the link between the two helicases and this cancer, we analysed the expression of *DDX17* and *DDX5* genes in the clinically annotated SEQC neuroblastoma cohort (Zhang et al., 2015). We found that a high expression of *DDX17* was significantly correlated with a better survival probability of patients (Fig. 3C, left panel). This was true also when the Kaplan-Meyer analysis was performed only on the most aggressive stage 4 patients (according to the International Neuroblastoma Staging System INSS) (Fig. 3C, right panel). Specifically, *DDX17* expression level is significantly lower in stage 4 tumours than in any other neuroblastoma stage (Fig. 3D), and is also significantly associated with high-risk tumours (Fig. 3E). Finally, we monitored *DDX17* expression according to the amplification of the *MYCN* oncogene, which is a well-established marker of poor prognostic in neuroblastoma (Otte et al., 2020), and we found a negative association between these two parameters (Fig. 3F). The same analysis showed a more contrasting result for the *DDX5* gene, but a low *DDX5* expression was also found to correlate with a lower survival probability and with *MYCN* amplification (Supp. Fig. 3).

In conclusion, those results established a link between the expression of *DDX17*, and to a lower extent of *DDX5*, and the severity of neuroblastoma. This raised the interesting possibility that a suboptimal expression of DDX17 and DDX5, which leads to transcription termination defects of hundreds of genes in cultured cells, could favour the abnormal expression of a subset of chimeric transcripts in aggressive neuroblastomas.

### MYCN increases transcriptional readthrough and chimeric transcript formation

The negative link between *MYCN* amplification and a lower expression of both *DDX17* and *DDX5* genes suggested that MYCN could control the expression of both helicases, but also that chimeric transcript expression could be increased upon MYCN overexpression. To test this double hypothesis, we used two different neuroblastoma cell lines with and without an amplification of *MYCN* to modulate the expression of the oncogene.

First, we transfected SH-SY5Y cells (no *MYCN* amplification) with increasing amounts of an HA-tagged MYCN-encoding plasmid. MYCN protein was barely detectable in control cells and strongly induced upon transfection, but this did not result in any significant change in DDX17 and DDX5 protein levels (Fig. 4A). Under these conditions, we observed an increased production of chimeric transcripts (Fig. 4B, Supp. Fig. 4). MYCN also induced transcription readthrough beyond the 3’ end of genes that we had previously identified as regulated by DDX17 and DDX5 (Terrone et al., 2022) (Fig. 4B right panel, Supp. Fig. 4).

**Figure 4.**
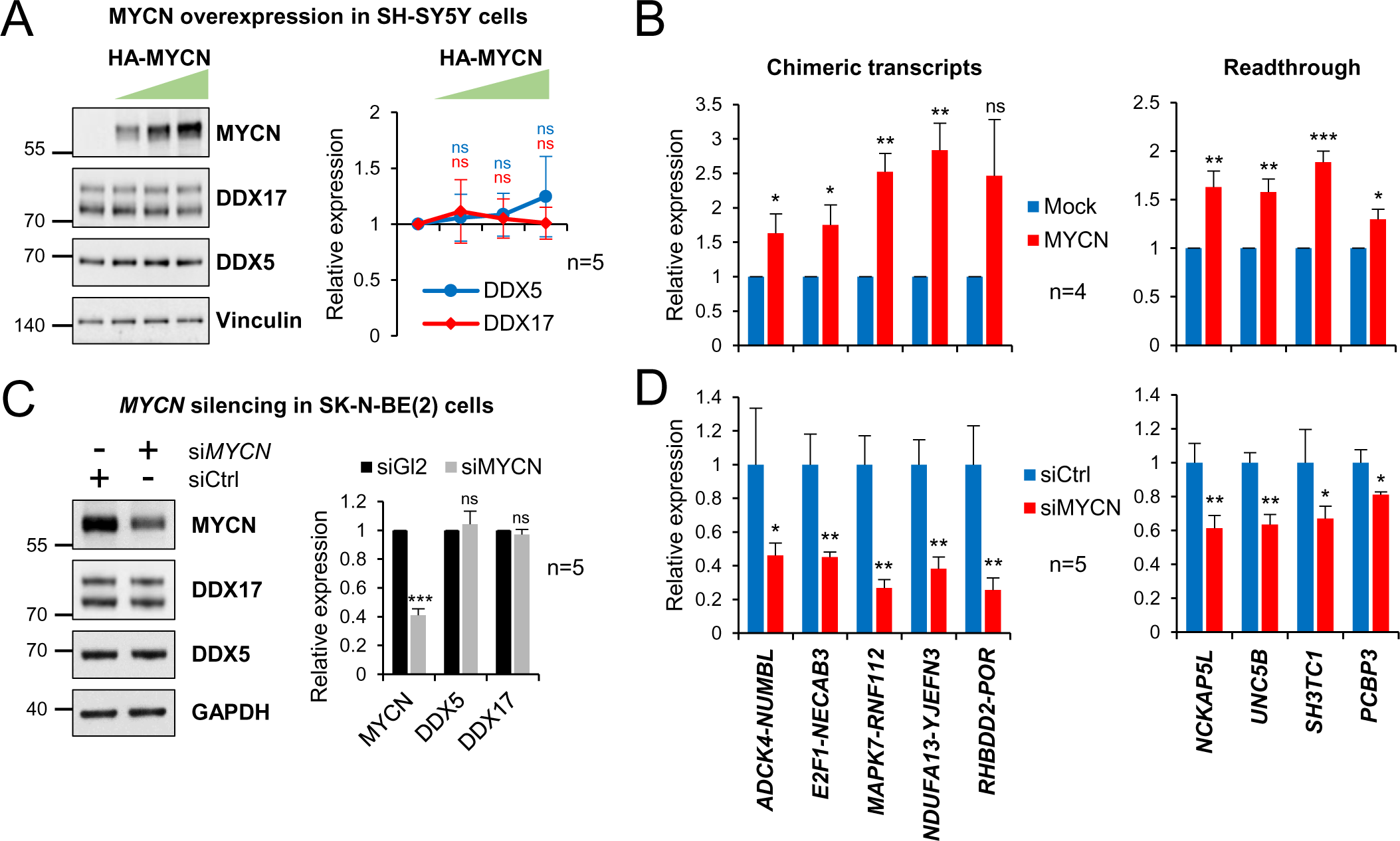
MYCN overexpression does not impact the expression of DDX17 and DDX5 but it induces the formation of chimeric transcripts. **A.** Western-blot showing DDX17 and DDX5 proteins levels upon increasing overexpression of MYCN in SH-SY5Y cells. Quantification of the western-blot (normalized to vinculin) is shown on the right. **B.** Expression of chimeric and readthrough transcripts upon MYCN overexpression. Details are as in Figure 1D (see also Supp. Fig. 4). Ratio paired *t*-test (*P-val<0.05 ; **P-val<0.01 ; ***P-val<0.001). **C.** Western-blot showing DDX17 and DDX5 proteins levels upon MYCN depletion in SK-N-BE(2) cells. Quantification of the western-blot (normalized to GAPDH) is shown on the right. **D.** Expression of chimeric and readthrough transcripts upon MYCN depletion. Unpaired Mann-Whitney test (*P-val<0.05 ; **P-val<0.01).

In parallel, we carried out the opposite experiment and downregulated MYCN expression in the *MYCN*-amplified SK-N-BE(2) cell line, using a mixture of specific siRNAs which reduced MYCN protein level by nearly 60%. Under these conditions, neither DDX17 nor DDX5 protein level was altered (Fig. 4C). However, we observed a significant reduction in the expression of previously tested chimeric transcripts, as well as reduced transcriptional readthrough (Fig. 4D, Supp. Fig. 4). Because steady-state levels of the corresponding canonical transcripts remained largely unaffected (Supp. Fig.4), our results suggested that variations in MYCN expression altered transcription termination and as a consequence, chimeric RNA production, in a manner that is distinct from its promoter-associated activity.

### MYCN binds near the 3’ end of DDX17/DDX5-regulated genes and interacts with DDX17

To test this hypothesis of a direct effect of MYCN on transcription termination, we re-analysed previously published ChIP-seq datasets performed in several neuroblastoma cell lines (Buchel et al., 2017, Upton et al., 2020, Zeid et al., 2018). We found that the 3’ region of genes displaying transcriptional readthrough or chimeric RNA production upon DDX17/DDX5 knock-down or MYCN overexpression generally exhibited at least one MYCN peak, often detected across several cell lines (Fig. 5A and Supp. Fig. 5). A global analysis of all DDX17/DDX5-regulated genes confirmed this observation, as the chromatin region overlapping their terminal exon was significantly enriched in MYCN binding sites, compared to non-regulated genes (Fig. 5B). We then performed ChIP-qPCR in SK-N-BE(2) cells and validated the binding of MYCN near the 3’ end of our model genes (Fig. 5C). Together with results of Figure 4 showing that MYCN does not affect DDX17 and DDX5 expression, these experiments strongly suggested that the effect of MYCN on transcription termination is direct.

**Figure 5.**
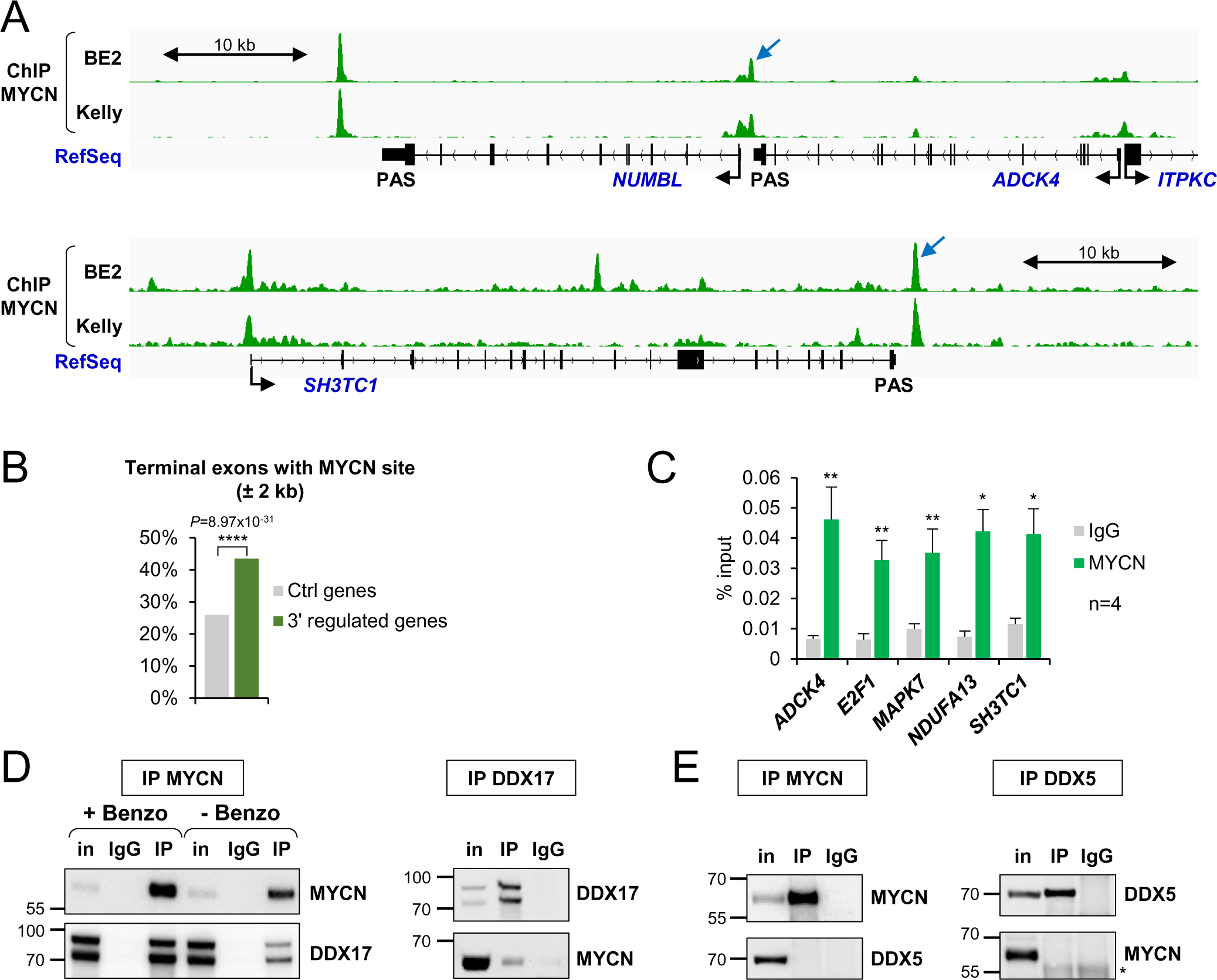
MYCN binds to the 3’ end of DDX17/DDX5-regulated genes and interacts with DDX17. **A.** Typical examples of DDX17/DDX5-regulated genes displaying MYCN binding near their 3’ end. **B.** Genome wide analysis of MYCN binding near terminal exons of DDX17/DDX5-regulated genes and control genes in neuroblastoma cell lines. Tukey’s test. **C.** ChIP-qPCR analysis of MYCN binding near the 3’ end of DDX17/DDX5-regulated genes. Paired *t*-test (*P-val<0.05 ; **P-val<0.01). **D.** Co-immunoprecipitation of endogenous MYCN and DDX17 in SK-N-BE(2) cells, in the presence or absence of benzonase, as indicated. **E.** Co-immunoprecipitation of endogenous MYCN and DDX5 in SK-N-BE(2) cells.

We next hypothesized that through its binding to chromatin at the 3’ end of DDX17/DDX5 target genes, MYCN could interfere with the function of helicases. Therefore, we tested whether MYCN could interact with DDX17 and DDX5 by carrying out co-immunoprecipitation (co-IP) assays in SK-N-BE(2) cells. As shown in Fig. 5D, endogenous DDX17 and MYCN efficiently co-immunoprecipitated, whatever protein was targeted. The co-IP was even enhanced when extracts were treated with benzonase, excluding the contribution of nucleic acids and suggesting a direct interaction between the two proteins. In contrast, no co-IP was detected between MYCN and DDX5 (Fig. 5E), which underlines the specificity of the MYCN-DDX17 interaction.

Altogether these results indicate that MYCN impacts transcription termination in a direct manner, possibly by inhibiting the normal function of DDX17.

### The forced recruitment of MYCN near the termination region is sufficient to induce the formation of chimeric transcripts

We sought to provide a direct proof that MYCN binding near the termination region of a gene is sufficient to promote transcriptional readthrough and the subsequent production of a chimeric transcript. We designed an experiment in SH-SY5Y cells (in which endogenous MYCN level is very low) in which we could force the recruitment of MYCN downstream of the PAS of the *MAPK7* gene. For this, we co-transfected cells with two plasmids, one expressing MYCN protein fused to a catalytically inactive "dead" Cas9 protein (dCas9-MYCN) and the other expressing specific RNA guides to target dCas9-MYCN downstream of the *MAPK7* gene (Fig. 6A). We verified by ChIP-qPCR that dCas9-MYCN was indeed bound at its target site upon transfection of RNA guides (Fig. 6B). A plasmid expressing dCas9-GFP fusion protein was used as a control to rule out a possible non-specific action of the dCas9 fusion protein on transcription through steric hindrance.

**Figure 6.**
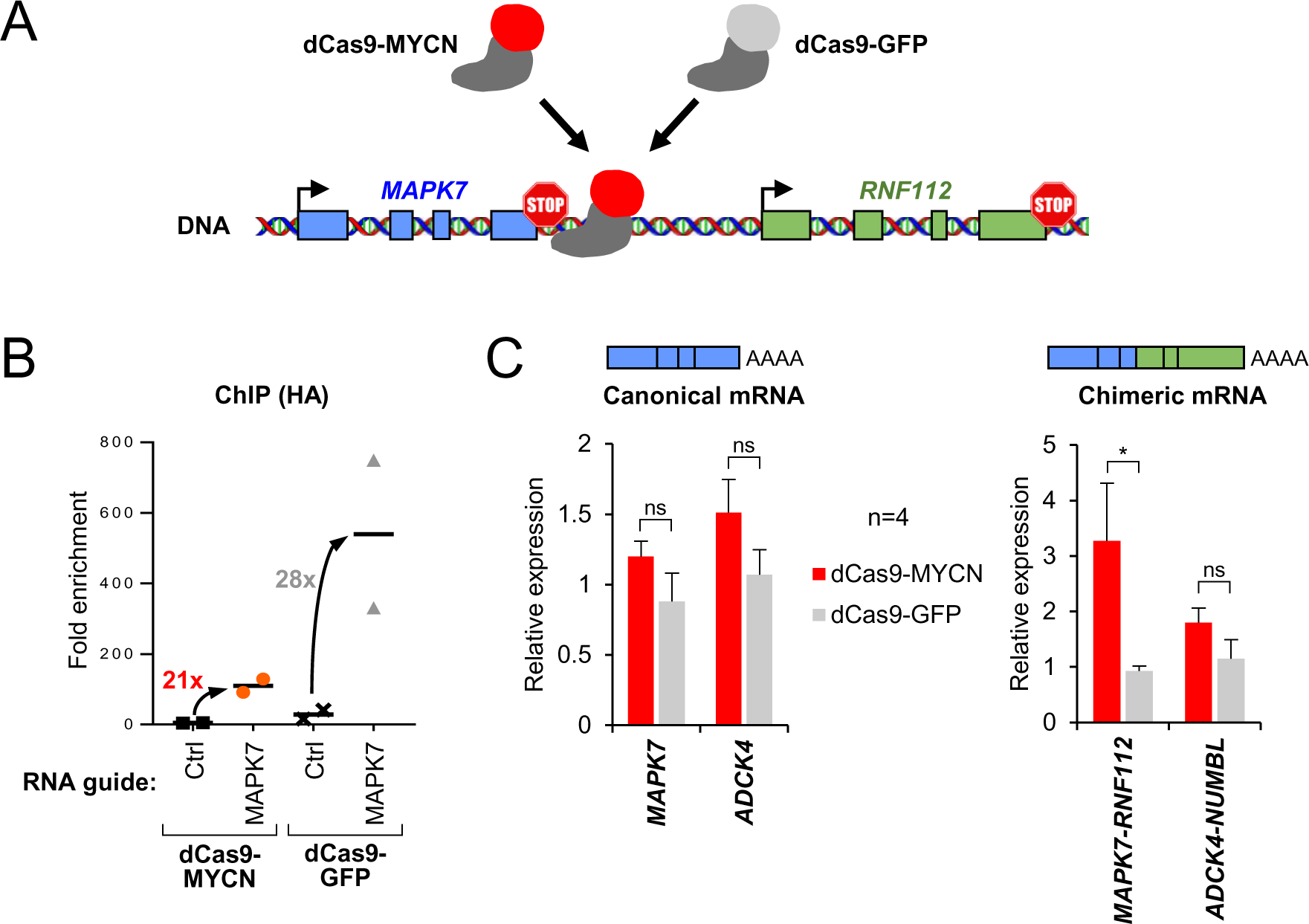
The forced recruitment of MYCN near the termination region is sufficient to induce the formation of chimeric transcripts. **A.** Strategy for the recruitment of MYCN (fused to a catalytically inactive dCas9) at the 3’ end of the *MAPK7* gene. **B.** ChIP-qPCR analysis of dCas9-MYCN and dCas9-GFP binding near the 3’ end of *MAPK7*. Each ChIP signal was normalised to its respective control (ChIP with IgG). **C.** Quantification of canonical *MAPK7* transcript and chimeric *MAPK7-RNF112* transcript in cells transfected with dCas9-MYCN or dCas9-GFP, relative to transfection without specific gRNA. Paired *t*-test (*P-val<0.05 ; **P-val<0.01).

We then quantified the expression of the *MAPK7* gene, which remained unaffected by the recruitment of either dCas9-MYCN or dCas9-GFP, as did the control *ADCK4* gene (Fig. 6B, left panel). In contrast, the chimeric *MAPK7-RNF112* chimer was induced only upon the binding of the dCas9-MYCN between the two genes while the *ADCK4-NUMBL* chimer remained unchanged (Fig. 6B, right panel).

This experiment demonstrated that the binding of MYCN near the termination site can inhibit the normal termination process and promote the formation of a chimeric transcript.

## Discussion

In this study, we show that the transcription readthrough induced by depletion of DDX17 and DDX5 helicases also leads to the production of chimeric transcripts in neuroblastoma cells. Among the most significantly induced chimeric transcripts, 27% were previously found in neuroblastoma tumors, and not in non-tumoural adrenal gland tissue (Shi et al., 2021). This suggests that this subset of chimeric transcripts might be specific of neuroblastomas and could potentially be used as biomarkers for this cancer. Further studies will be required to determine their level of specificity, as chimeric transcripts have been identified in many types of cancers (Dorney et al., 2023, Sun and Li, 2022).

If production of chimeric transcript in healthy cells is considered as a possible mechanism to increase protein diversity (Parra et al., 2006), their abnormal expression in cancer cells can have several consequence. First, a fraction of chimeric transcripts can retain a functional ORF allowing the translation of chimeric or fusion proteins (Varley et al., 2014, Yun et al., 2014, Han et al., 2017, Egashira et al., 2019), and we validated the production of CTSD-IFITM10 protein upon DDX7/DDX5 depletion. Chimeric proteins can generate highly immunogenic neo-antigens, which may be unique to the tumor and could be targeted by the immune system (Weber et al., 2022). A recent study identified a novel chimeric transcript produced by gene fusion and demonstrated that it produces a neo-antigen that can specifically elicit a host cytotoxic T-cell response in metastatic head and neck cancer (Yang et al., 2019). Chimeric proteins produced from readthrough induced transcripts may be much less abundant than those produced from a gene fusion event, but depending on their expression level, they could alter the protein interaction network and activate cell growth pathways by acting as oncoproteins, or alter the cell response to antitumour treatments. Second, even if not translated, chimeric transcripts can act as functional long noncoding RNAs (lncRNA). For example, *SLC45A3-ELK4* chimeric transcripts act as lncRNAs and regulate cell growth in prostate cancer, and its higher expression level has been associated with disease progression and metastasis (Qin et al., 2017, Zhang et al., 2012). Future analyses will determine whether the chimeric transcripts we have identified have protumoral properties or can influence tumour response to treatment.

Our results suggest that the production of chimeric transcripts in neuroblastomas could be favoured by a reduced expression of *DDX17* and *DDX5*, especially in high-risk tumours. Indeed, a low expression of both helicases is significantly associated with higher risk and shorter patient survival. Note that our results differ from a previous report showing a link between high *DDX5* expression and poor survival of patients with neuroblastoma (Zhao et al., 2020). However, we used a much larger and more detailed cohort of tumours (n=498 versus n=42), which allowed us to stratify the expression of helicases across all neuroblastoma stages. Importantly, the fact that high expression of *DDX17* and *DDX5* is associated with better survival when looking at stage 4 patients only, reduces the possibility that it is directly and solely linked to *MYCN* amplification, which is a less predominant risk factor in this class of tumours. It will be important to confirm our results at the protein level, and to determine whether variations in the expression of both helicases contribute to the mechanism of oncogenesis in neuroblastoma.

Interestingly, neuroblastoma is only one of few examples of cancers in which a high expression of *DDX17* and/or *DDX5* is associated with good patient survival, among many studies describing their positive contribution to major cancer signaling pathways and cancer development (Fuller-Pace and Moore, 2011, Xu et al., 2022). This suggests that the two helicases play contrasted roles depending probably on the cancer type and on the context in which they are studied. Beside neuroblastoma, high *DDX17* expression and high *DDX5* expression are respectively associated with a good prognosis in breast cancer (Wortham et al., 2009) and lung adenocarcinoma (Zhou et al., 2024), and with pancreatic ductal adenocarcinoma (Morimachi et al., 2021) and hepatocellular carcinoma (Zhang et al., 2016a, Zhang et al., 2019). In addition to their multiple roles in controlling gene expression and RNA processing, for example by repressing readthrough transcription, DDX5 and DDX17 are also involved in genomic stability and DNA repair (Bader et al., 2022, Cargill et al., 2021), which may also explain how their low expression can impact high-risk tumours.

We show that MYCN, a transcription factor with a major oncogenic effect in neuroblastoma, promotes transcriptional readthrough and the production of chimeric transcripts, without altering the expression of either helicase. Our results indicate that this new function of the oncoprotein is direct, yet the mechanism by which it inhibits termination is still unclear. MYCN co-immunoprecipitates with DDX17 and binds preferentially downstream of DDX17/DDX5-regulated genes, suggesting that it may interfere with the activity of the helicases. We showed previously that those genes present a looped 3D conformation and that their transcription termination of is also controlled by CTCF (Terrone et al., 2022), so it will be interesting to test whether MYCN overexpression impacts gene looping and CTCF binding. Supporting this idea, MYCN and CTCF bind to hundreds of overlapping intergenic sites (Lee et al., 2012, Buchel et al., 2017), and MYC was recently shown to affect CTCF binding (Wei et al., 2023). Another hypothesis is that MYCN could prevent termination at the end of the gene by activating RNAPII, as MYCN or MYC do to induce promoter escape and productive elongation (Baluapuri et al., 2019, Herold et al., 2019, Papadopoulos et al., 2022). Alternatively, MYCN interacts and recruits multiple chromatin remodeling complexes, whose ectopic recruitment to the 3’ end of gene may alter the local chromatin state and affect RNAPII kinetics and interactome and disturb the chain of events leading to termination.

Future work will help to determine which of these non-mutually exclusive hypotheses is correct, or if another mechanism is involved. Nevertheless, our study is a new illustration of the diversity of actions that a transcription factor can have upon its binding to non-promoter sites. To summarize, our results show that MYCN overexpression or a suboptimal expression of either DDX17 or DDX5, two parameters associated to poor survival of neuroblastoma patients, induce transcription readthrough of a subset of common genes and promote the production of chimeric transcripts. Understanding the interplay between these factors is of great importance to appreciate their impact on tumorigenesis.

## Materials and Methods

### Plasmids and cell culture

Human SH-SY5Y and SK-N-BE(2) cells (ECACC) were grown and transfected essentially as described previously (Lambert et al., 2018). For standard silencing experiments, 20 nM of siRNA (sequences in Supp. Table 2) were mixed with Lipofectamine™ RNAiMax (ThermoFisher Scientific) following the manufacturer’s instructions and cells were harvested 48 h after transfection. For MYCN overexpression, cells were plated in 6-well-plates to reach 70% confluence, and then transfected with 0.5 to 1 μg pCDNA3-HA-hMYCN (Addgene #74163) using JetPrime® (Polyplus Transfection) following the manufacturer’s instructions. Cells were harvested 48 h after transfection.

For dCas9-MYCN experiments, the hMYCN cDNA was subcloned into the dCas9-empty-GFP plasmid (a gift from Reini F. Luco, Institut Curie, Orsay, France). This cloning step was done by RD Biotech. We cloned the sequences corresponding to the two RNA guides into the *Bsm*BI site of the CRIZI plasmid (provided by Philippe Mangeot, CIRI, Lyon, France). SH-SY5Y cells were plated in 6-well-plates to reach 70% confluence and transfected with 1 μg of dCas9-HA-MYCN plasmid and 1 μg of sgRNA-containing plasmid (500 ng of each guide, sequences in Supp. Table 2) using jetPRIME (PolyPlus Transfection).

### Western Blot and co-immunoprecipitation

Protein extraction and western blotting were carried out as previously described (Dardenne et al., 2014). Primary antibodies used for western-blotting were: DDX5 (ab10261, Abcam), DDX17 (ab24601, Abcam), GAPDH (sc-32233, Santa Cruz Biotechnology), MYCN (sc-53993, Santa Cruz Biotechnology), CTSD (21327-1-AP, Proteintech). For co-immunoprecipitation, SH-SY5Y cells were harvested and gently lysed for 5 min on ice in a buffer containing 10 mM Tris-HCl pH 8.0, 140 mM NaCl, 1.5 mM MgCL_2_, 10 mM EDTA, 0.5% NP40, completed with protease and phosphatase inhibitors (Roche #11697498001 and #5892970001), to isolate the nuclei from the cytoplasm. After centrifugation, nuclei were lysed in IP buffer (20 mM Tris-HCl pH 7.5, 150 mM NaCl, 2 mM EDTA, 1% NP40, 10% glycerol and protease/phosphatase inhibitors) for 30 min at 4°C under constant mixing. The nuclear lysate was centrifuged for 15 min to remove cell debris and soluble proteins were quantified by BCA (ThermoFisher). The lysate was pre-cleared with 30 μg of Dynabeads Protein G/A (ThermoFisher) for 30 min under rotating mixing, and then split in aliquots of 1.5 mg proteins for each assay. Each fraction received 5 μg of antibody and the incubation was left overnight at 4°C under rotating mixing. The following antibodies were used for IP: rabbit anti-DDX17 (19910-1-AP, Proteintech) or control rabbit IgG (ThermoFisher), goat anti-DDX5 (ab10261, Abcam) or control goat IgG (Santa Cruz Biotechnology), and mouse anti-MYCN (sc-53993, Santa Cruz Biotechnology) or control mouse IgG. The next day, the different lysate/antibody mixtures were divided, treated or not with 250 U/ml benzonase (Merck-Millipore) for 30 min at 37°C, and then incubated with 50 μg Dynabeads Protein G/A (ThermoFisher) blocked with bovine serum albumin, for 4 h at 4°C under rotating mixing. Bead were then washed 5 times with IP buffer. Elution was performed by boiling for 5 min in SDS-PAGE loading buffer prior to analysis by western-blotting.

### RNA extraction and real-time quantitative PCR

Total RNAs were isolated using TriPure Isolation Reagent (Roche). For reverse transcription, 2 μg of purified RNAs were treated with Dnase I (ThermoFisher) and retrotranscribed using Maxima reverse transcriptase (ThermoFisher), as recommended by the supplier. Potential genomic DNA contamination was systematically checked by performing negative RT controls in the absence of enzyme and by including controls with water instead of cDNA in qPCR assays. PCR reactions were carried out described previously (Lambert et al., 2018). For qPCR analyses, the specificity and linear efficiency of each primer set (sequences are in Supp. Table 2) was first verified by establishing a standard expression curve with various amounts of human genomic DNA or cDNA. qPCR reactions were carried out on 0.625 ng of cDNA using a LightCycler 480 System (Roche), with the SYBR® Premix Ex Taq (Tli RNaseH Plus, Takara), under conditions recommended by the manufacturer. Melting curves were controlled to rule out the existence of non-specific products. Relative DNA levels were calculated using the 2-ΔΔCt method (using the average Ct obtained from technical duplicates or triplicates) and were normalized to the expression of *GAPDH* RNA.

### Chromatin immunoprecipitation

A total of 10^7^ cells were crosslinked with 1% formaldehyde for 10 min at room temperature. Crosslinking was quenched by addition of 0.125 M glycin. Nuclei were isolated by sonication using a Covaris S220 (2 min, Peak Power: 75; Duty Factor: 2; Cycles/burst: 200), pelleted by centrifugation at 1000 g for 5 min at 4°C, washed once with FL buffer (5 mM HEPES pH 8.0, 85 mM KCl, 0.5% NP40) and resuspended in 1 ml shearing buffer (10 mM Tris-HCl pH 8.0, 1 mM EDTA, 0.1% SDS). Chromatin was sheared in order to obtain fragments ranging from 200 to 800 bp using Covaris S220 (20 min, Peak Power: 140; Duty Factor: 5; Cycles/burst: 200). Chromatin was next immunoprecipitated overnight at 4°C with 4 μg of mouse anti-MYCN II antibody (sc53993, Santa Cruz Biotechnology) or an equivalent amount of the corresponding IgG Isotype control (ThermoFisher) and 30 μl of Dynabeads® Protein A/G (ThermoFisher). Complexes were washed with 4 different buffers: Low salt buffer (20 mM Tris-HCl pH 8.0, 150 mM NaCl, 1% Triton X-100, 0.1% SDS, 2 mM EDTA), High salt buffer (20 mM Tris-HCl pH 8.0, 500 mM NaCl, 1% Triton X-100, 0.1% SDS, 2 mM EDTA), Low LiCl buffer (10 mM Tris-HCl pH 8.0, 0.5 M LiCl, 1% NaDoc, 1% NP40), Tris/EDTA (10 mM Tris-HCl pH 8.0, 1 mM EDTA), and were eluted in Elution buffer (200 mM NaCl, 1% SDS, 0.1 M NaHCO_3_, 20 μg Proteinase K) overnight at 65°C. The immunoprecipitated chromatin was purified by phenol-chloroform extraction and ethanol precipitation, and analysed by qPCR. Values were normalized to the signal obtained for the immunoprecipitation with control IgG.

### *In silico* prediction of chimeric RNAs from RNA-seq data

The raw RNA-seq data were described previously (Terrone et al., 2022) and are accessible from the Gene Expression Omnibus repository (accession number GSE183205). Raw reads were pre-processed using fastp (v0.23.2) (Chen et al., 2018) and mapped to the human reference genome (hg19, GRCh37.87) using STAR (v2.7.8a) (Dobin et al., 2013). Mapped reads were filtered using samtools (v1.11) (Danecek et al., 2021). Gene fusions were detected using Arriba (v2.3.0) (Uhrig et al., 2021) and results were parsed using an R script. Split and discordant reads identified by Arriba were counted for each fusion and compared to control condition. Differentially fused genes (DFG) were analyzed using the DESeq2 package (v1.40.1) (Love et al., 2014). The complete pipeline is available in Nextflow (Di Tommaso et al., 2017) at https://gitbio.ens-lyon.fr/LBMC/regards/nextflow/-/blob/master/src/arriba_fusion.nf.

### Statistical analysis of *DDX17* and *DDX5* expression in neuroblastoma tumours

To test the expression of *DDX17* and *DDX5* in neuroblastomas, we used the previously described cohort of 498 tumours (GEO accession number: GSE62564) (Zhang et al., 2016b), in which patients groups were defined according to the International Neuroblastoma Staging System (INSS). Survival curves were generated by the Kaplan-Meier method, and statistical analyses were carried out as described previously (Gibert et al., 2014, Jiang et al., 2021).

### Analysis of MYCN ChIP-seq data

To analyse the relative proximity of DDX17/DDX5-regulated terminal exons to MYCN binding sites, we first generated a BED file containing a merged list of MYCN peaks identified in several neuroblastoma cell lines (Buchel et al., 2017, Upton et al., 2020, Zeid et al., 2018) (Supp. Table 3), re-analysed as follows. Raw reads were pre-processed using fastp (v0.23.2) (Chen et al., 2018) and mapped to the human reference genome (hg19, GRCh37.87) using bowtie2 (v2.5.2) (Langmead et al., 2019). Mapped reads were filtered using samtools (v1.11) (Danecek et al., 2021) and then formatted using deepTools2 (v3.5.1) (Ramirez et al., 2016). Peak calling analysis was carried out with MACS (v3.0.0a6) (Zhang et al., 2008). Nearest exons from peak summits were identified using to BEDtools (v2.25.0) (Quinlan and Hall, 2010). The complete ChIP-seq pipeline is available in Nextflow (Di Tommaso et al., 2017) at https://gitbio.enslyon.fr/xgrand/ChIPster.

We next calculated the genomic distance (negative or positive for upstream and downstream peaks, respectively) between each exon from the FasterDB database (Mallinjoud et al., 2014) and the summit of the closest MYCN peak. Only genes having at least one internal exon were considered. We performed a logistic regression analysis to test if the 3’ terminal exons regulated by DDX5/DDX17 (n = 933) are closer to MYCN peaks than unregulated terminal exons. (n = 18,482). We modeled the proximity to a MYCN peak according to the two groups of exons using the glm function, with family = binomial (‘logit’) in R software. A MYCN peak was considered as close to an exon if its center is located within an exon or within an interval of 1 base to 2 kb upstream or downstream of the exon. To test the differences between the two groups of exons, a Tukey’s test was used (with R, emmeans function from emmeans library).

## Acknowledgments

We wish to thank Reini F. Luco (Institut Curie, Orsay, France) and Philippe Mangeot (CIRI, Lyon, France) for sharing reagents. We are grateful to LBMC members for fruitful discussions and advice. We acknowledge the CBPSMN (Centre Blaise Pascal de Simulation et de Modélisation Numérique) of the ENS-Lyon for computing resources. This work was supported by grants from Ligue contre le Cancer (Equipe labellisée), Fondation ARC, and Association Hubert Gouin “Enfance et Cancer”. V.C. was supported by a doctoral fellowship from Fondation pour la Recherche Médicale (FRM). J.V. received doctoral fellowships from the French Ministry of Research and Education and from the Ligue contre le Cancer.

## Author Contributions

All authors contributed to the study conception and design. V.C., J.V. and M.B. carried out experiments, collected and interpreted the data. X.G. and N.F. analysed RNA-seq and ChIP-seq datasets. N.R. and B.G. analysed the cohort of neuroblastoma tumours. The first draft of the manuscript was written by V.C. and C.F.B. and all authors commented, corrected and approved the manuscript.

