## Supplementary legends and figures for "The MYCN oncoprotein and helicases DDX17 and DDX5 have opposite effects on the production of chimeric transcripts in neuroblastoma cells": clerc_supp_biorxiv.pdf

#### Legends for Supplementary Figures

**Supp. Figure 1.** Typical examples of chimeric trancripts induced by DDX17 and DDX5 depletion. **A.** Screenshot from the Integrative Genome Viewer (IGV) browser showing the RNA-seq reads aligning to *CTSD* and *IFITM10* genes. The sequencing coverage of control (blue) and siDDX17/DDX5 (red) conditions shows a reduction of the steady-state expression the *CTSD* gene, yet the number of reads spanning the junction between exons of both genes is higher upon DDX17/DDX5 depletion. Most reads characterizing the chimeric *CTSD-IFITM10* transcripts link the one-to-the-last exon of *CTSD* to the second exon of *IFITM10*. **B.** Other IGV screenshots, in which only the sequencing coverage and splicing junctions are shown. Control and siDDX17/DDX5 conditions are in red and blue, respectively.

**Supp. Figure 2. A.** Validation of the expression of chimeric transcripts by RT-PCR. Expression of chimeric transcripts was monitored using primers spanning the chimeric junction and compared to the expression of the parental gene. Controls for reverse transcription (RNA in the RT) and PCR (no cDNA) are included. The asterisk on the *GAPDH* panel indicates a non-specific PCR product (dimers of primers). **B.** RT-qPCR quantification of the steady-state expression of canonical mRNAs corresponding to the chimeric transcripts validated in Fig. 1C. Two-tailed paired t-test (* *P*-val<0.05 ; ** *P*-val<0.01).

**Supp. Figure 3.** The expression of the *DDX5* gene is reduced in high risk neuroblastomas. **A.** Kaplan-Meier curves showing the overall survival of neuroblastoma patients, separated in 2 groups of high and low *DDX5* expression (relative to the median value of the group). Left diagram: all patients (n=498). Right diagram: stage 4 patients (n=183). The number of patients in each group and for each time point is indicated below. **B.** Box plot of *DDX5* expression relative to tumor stage (INSS classification). ANOVA corrected for multiple comparisons with Tukey's tests. Only the comparison between Stage 4 and other stages is shown, the other comparisons were not significant. **C.** Box plot of *DDX5* expression relative to low/high risk classification. Two-tailed *t*-test. **D.** Box plot of *DDX5* expression relative to amplified (A) or non amplied (NA) *MYCN* status. Two-tailed *t*-test.

**Supp. Figure 4. A.** RT-qPCR quantification of the steady-state expression of canonical mRNAs corresponding to the chimeric and readthrough transcripts analysed in Fig. 4B, upon MYCN overexpression. Ratio paired *t*-test (*P-val<0.05 ; **P-val<0.01 ; ***P-val<0.001). **B.** RT-qPCR quantification of the steady-state expression of canonical mRNAs corresponding to the chimeric and readthrough transcripts analysed in Fig. 4D, upon *MYCN* silencing. Unpaired Mann-Whitney test (*P-val<0.05 ; **P-val<0.01).

**Supp. Figure 5.** Examples of DDX17/DDX5-regulated genes displaying MYCN binding near their 3' end in 5 different neuroblastoma cell lines analysed by ChIP-seq. Peaks identified as significant are indicated with yellow arrows.

#### Legends for Supplementary Tables

**Supp. Table 1.** List of the 282 chimeric transcrits whose expression is changed upon siDDX17/DDX5 treatment, identified by the Arriba pipeline. The 97 most significant chimeric transcripts have their p-value cell highlighted in green. The last 2 columns report the chimeric transcripts identified by Shi *et al* in neuroblastoma tumours, with their number of occurences and corresponding frequency. The ID of these transcripts are highlighted in red, and the 25 most significant are in yellow characters. The second tab of the table compiles the merged list of genes whose termination is affected by DDX17/DDX5 depletion, generating readthrough transcription (our previous work, Terrone *et al*., 2022) and chimeric transcript formation (this study).

**Supp. Table 2.** List of primers and siRNAs used in this study.

**Supp. Table 3.** List of public ChIP-seq datasets used to analyse MYCN binding.
PMID 32286315: Upton K *et al.* (2020). Epigenomic profiling of neuroblastoma cell lines. Sci Data 7(1):116. doi: 10.1038/s41597-020-0458-y.
PMID 29262328: Büchel *et al.* (2017). Association with Aurora-A Controls N-MYC-Dependent Promoter Escape and Pause Release of RNA Polymerase II during the Cell Cycle. Cell Rep 21(12):3483-3497. doi: 10.1016/j.celrep.2017.11.090.
PMID 29379199: Zeid R *et al.* (2018). Enhancer invasion shapes MYCN-dependent transcriptional amplification in neuroblastoma. Nat Genet 50(4):515-523. doi: 10.1038/s41588-018-0044-9.

### Supp. Figure 1A

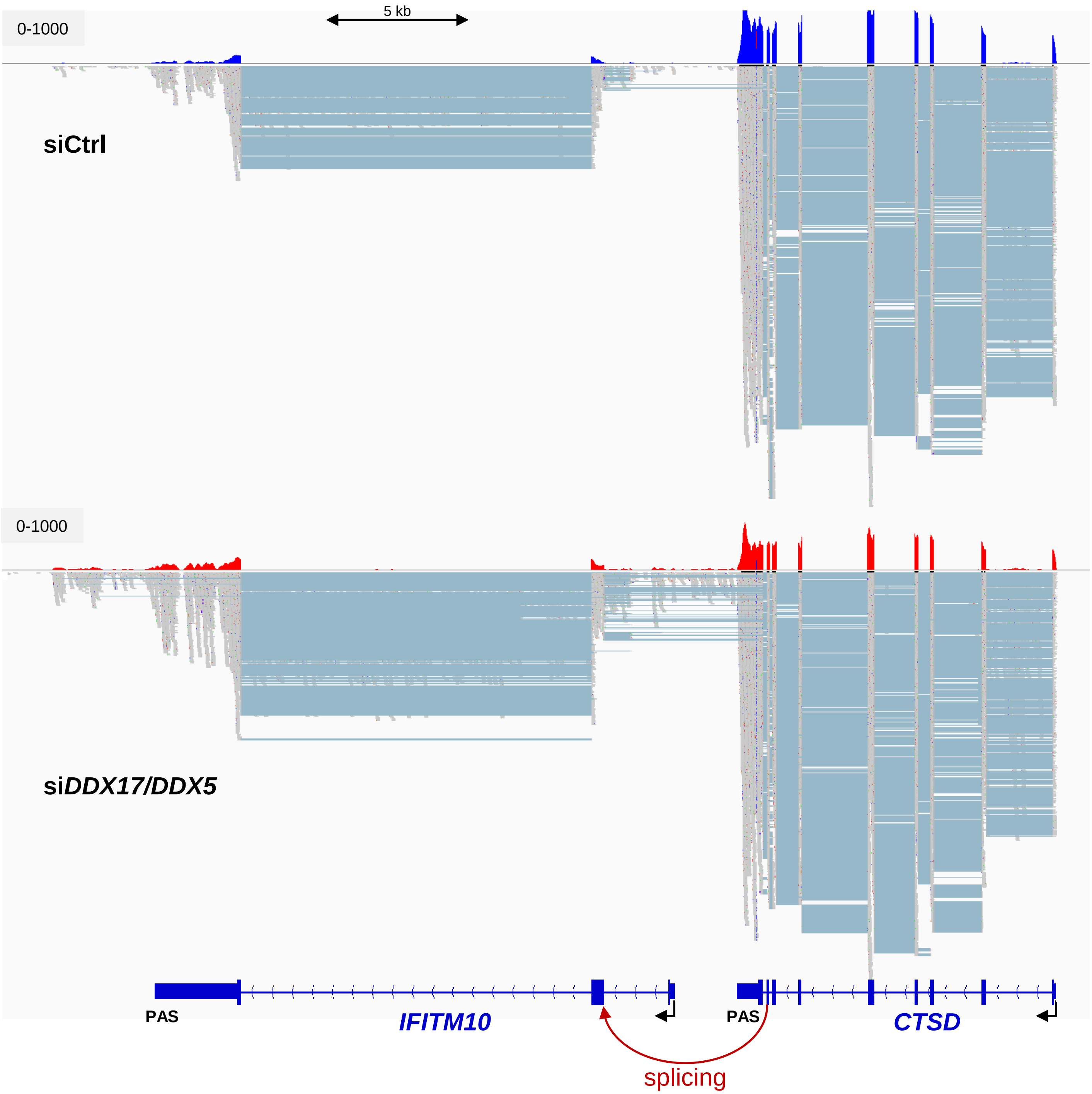

### Supp. Figure 1B

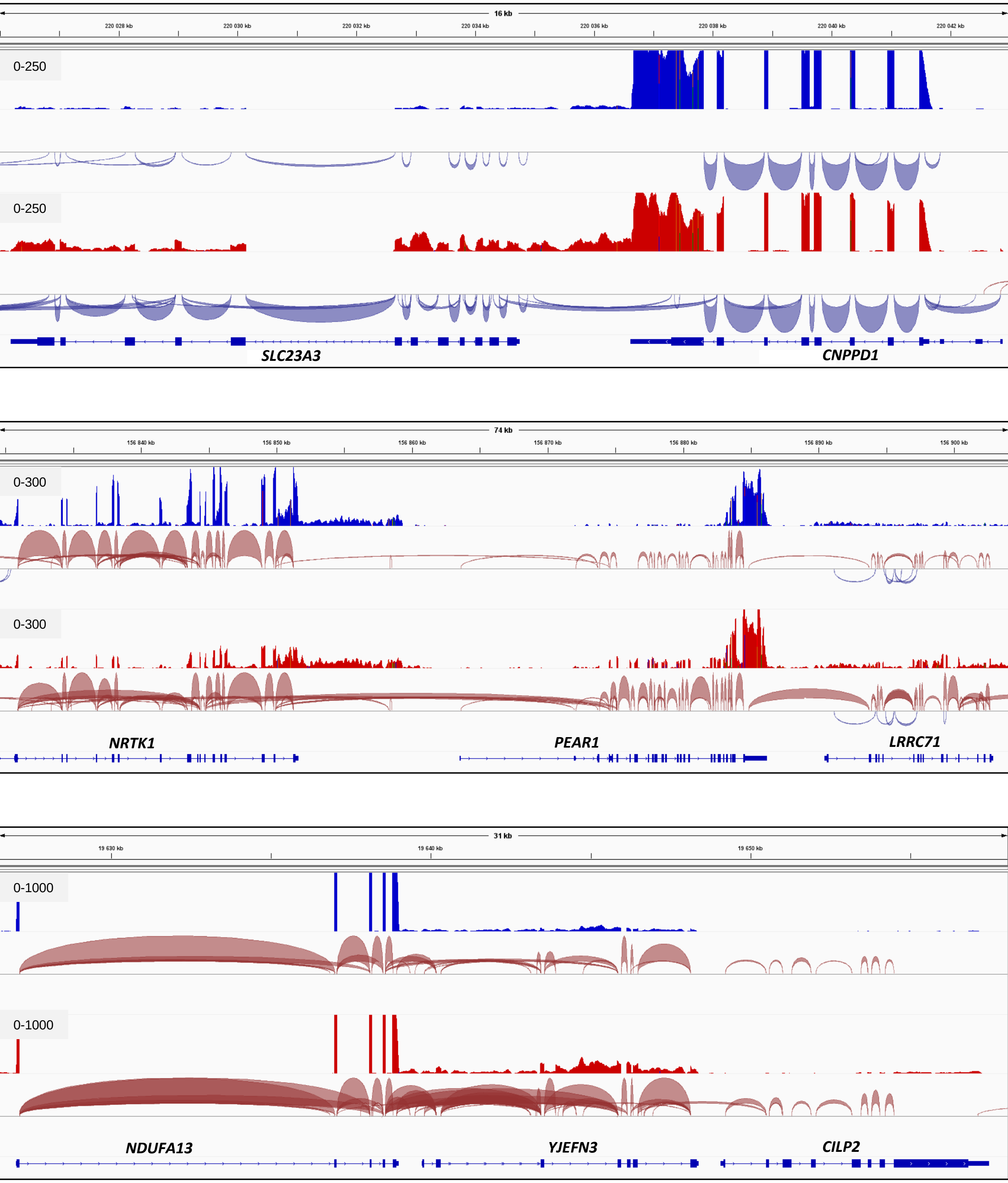

### Supp. Figure 1B

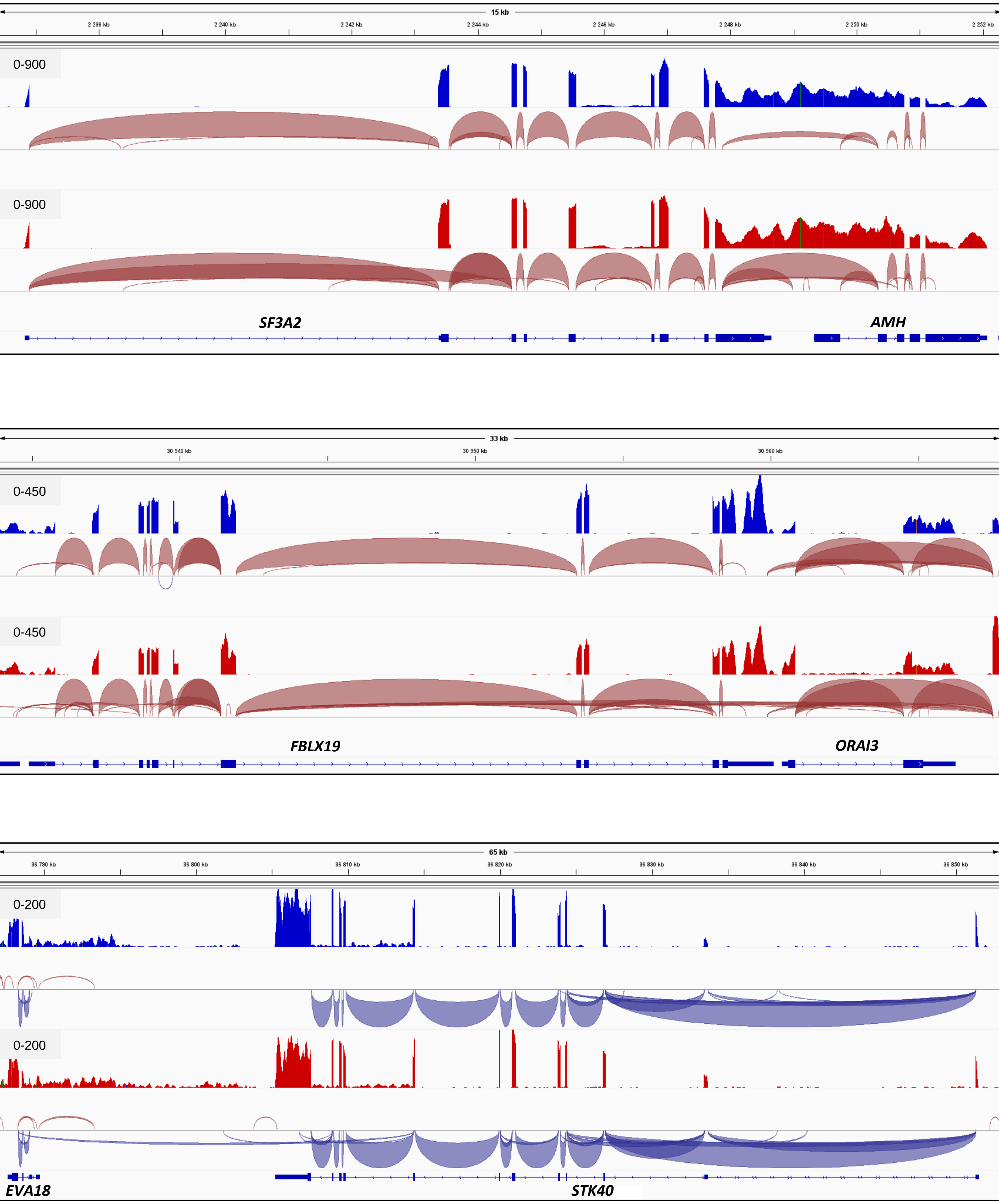

### Supp. Figure 1B

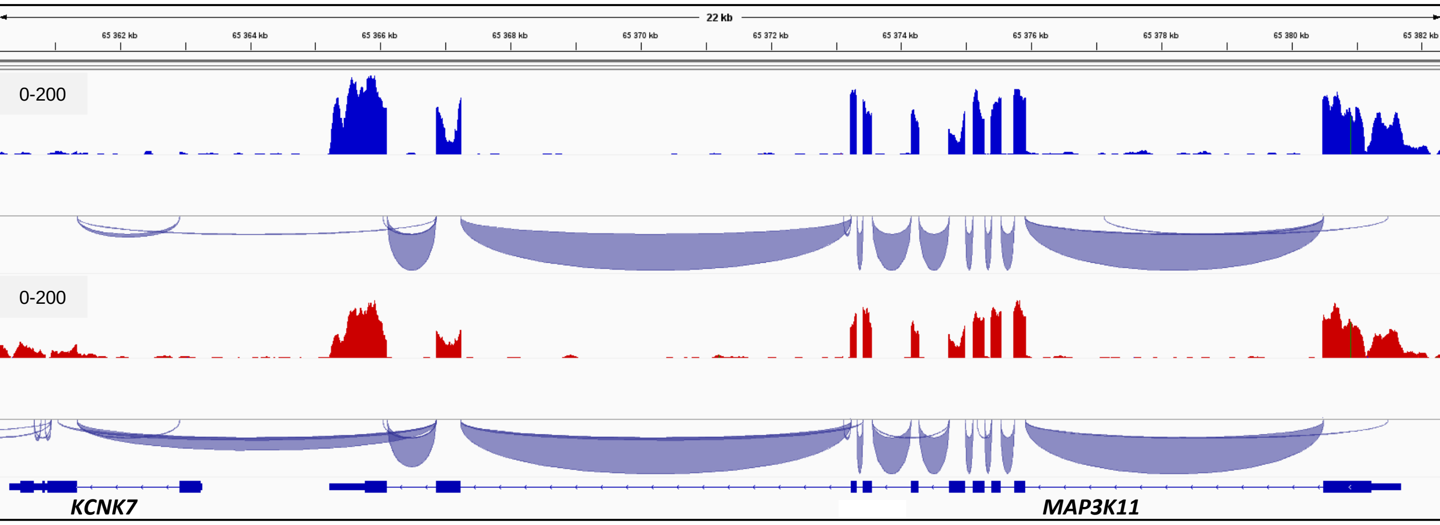


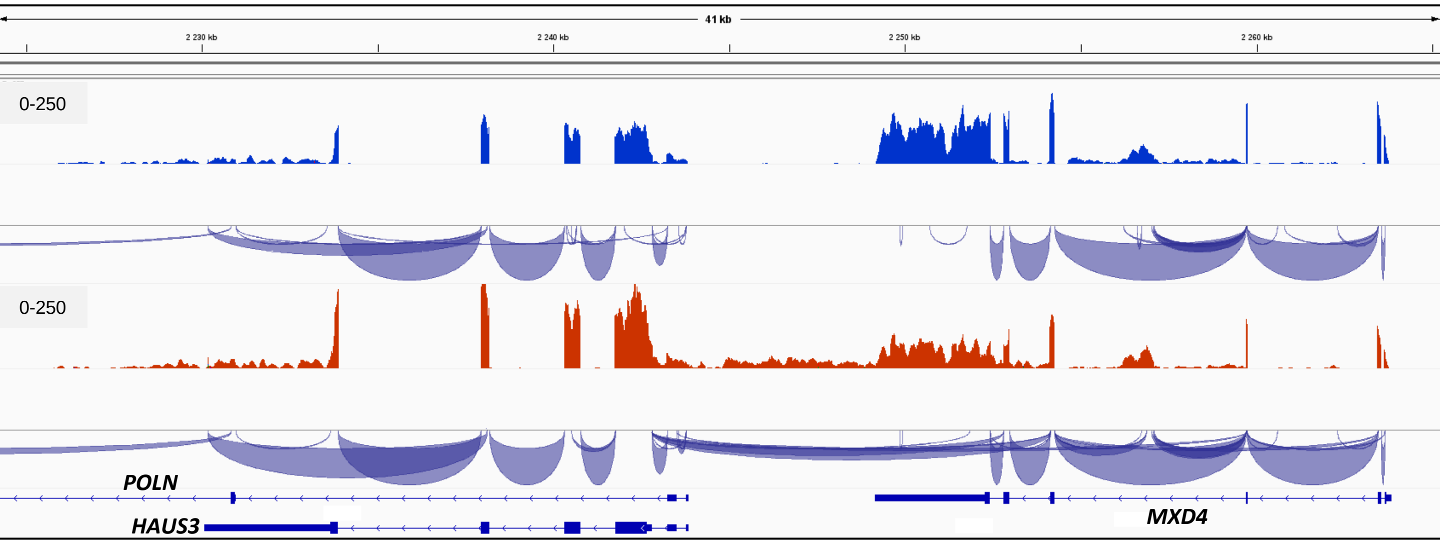

### Supp. Figure 2

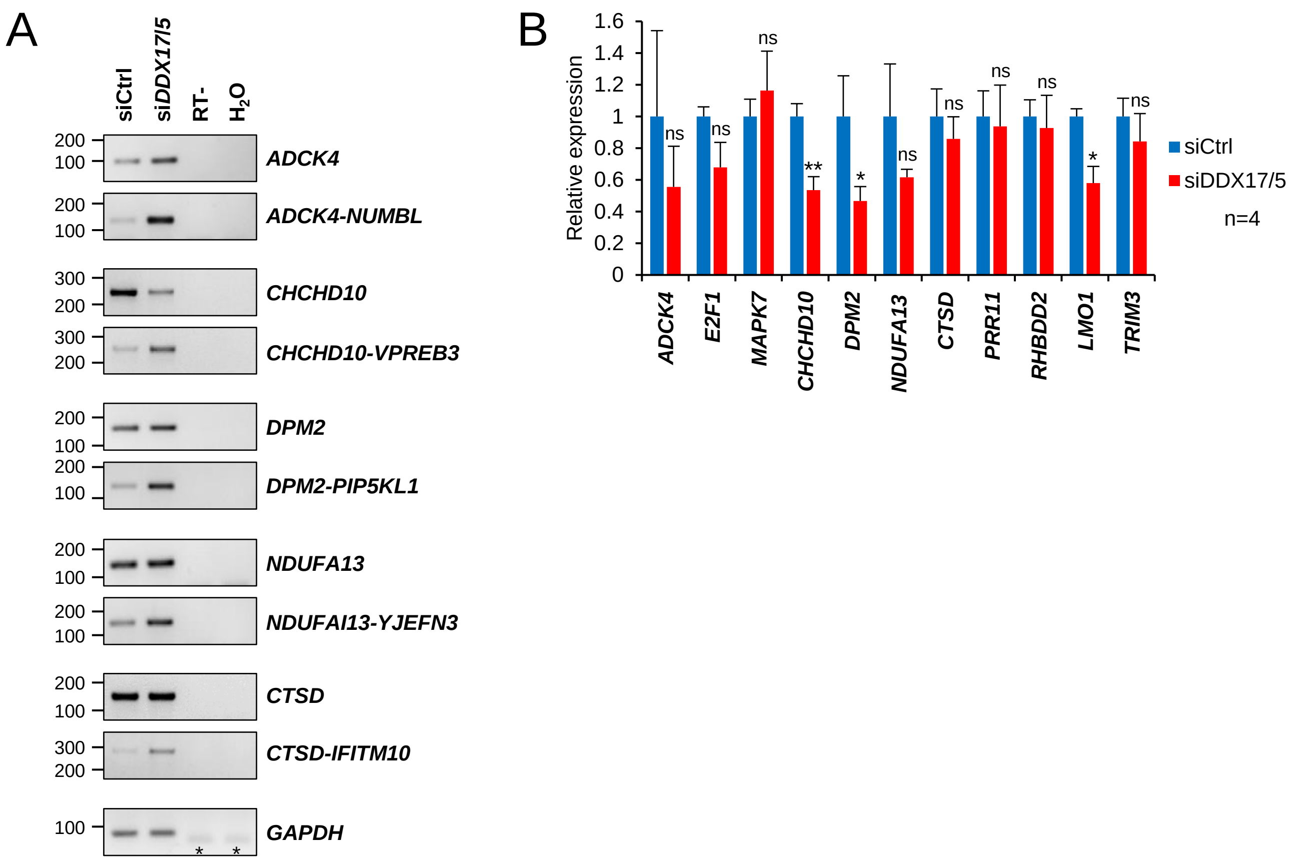

### Supp. Figure 3

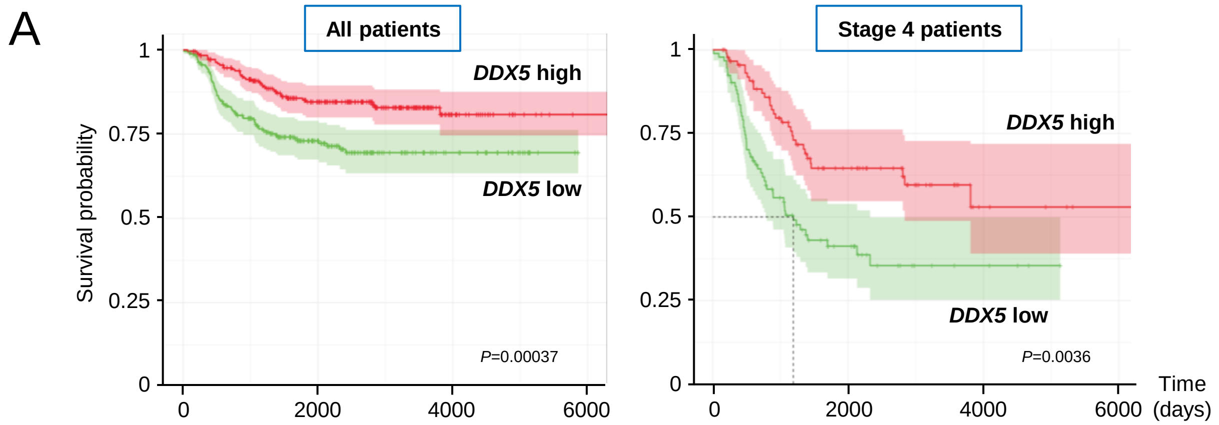


| | | | | | | | | |
|---|---|---|---|---|---|---|---|---|
| ***DDX5* low** | 249 | 103 | 25 | 0 | 92 | 20 | 4 | 0 |
| ***DDX* high** | 249 | 147 | 27 | 2 | 91 | 38 | 5 | 1 |

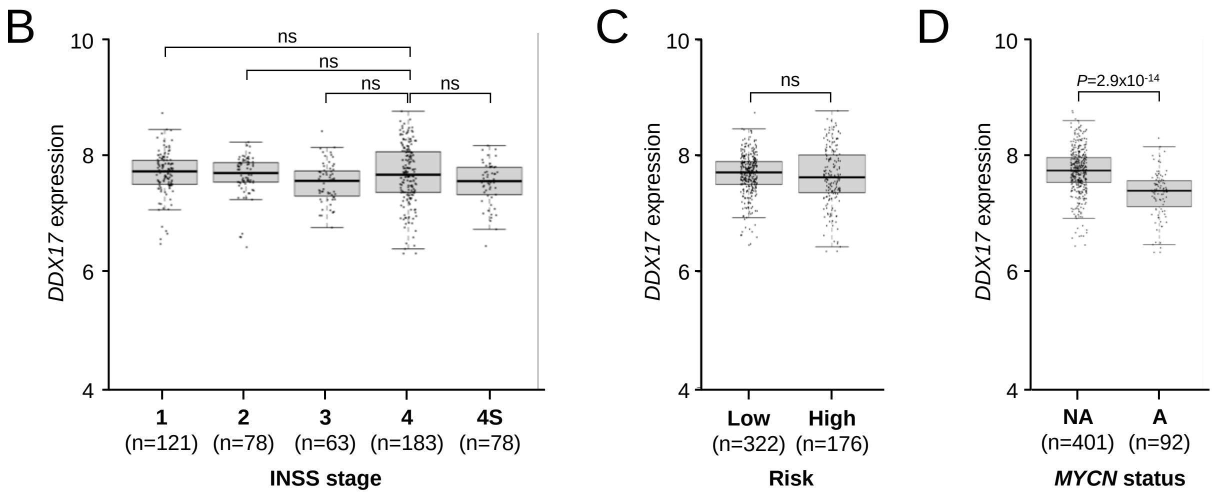

### Supp. Figure 4

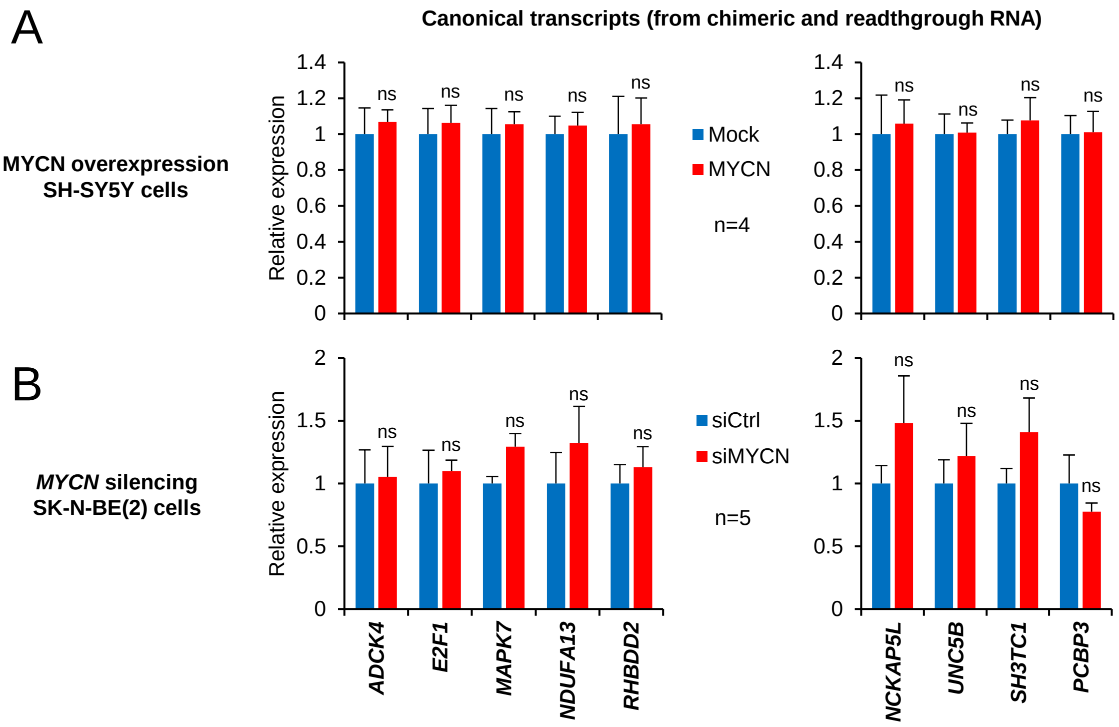

### Supp. Figure 5

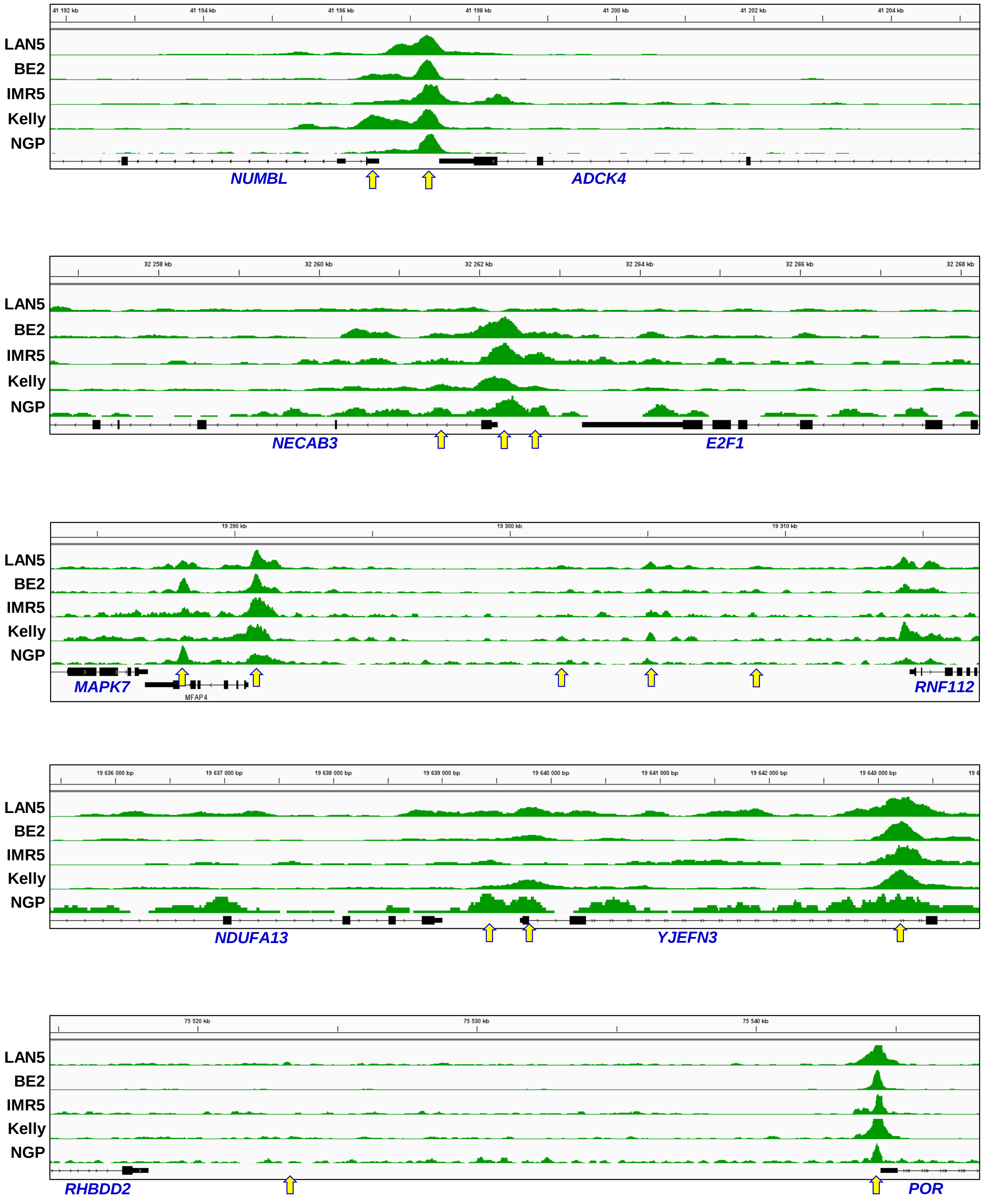

### Supp. Figure 5

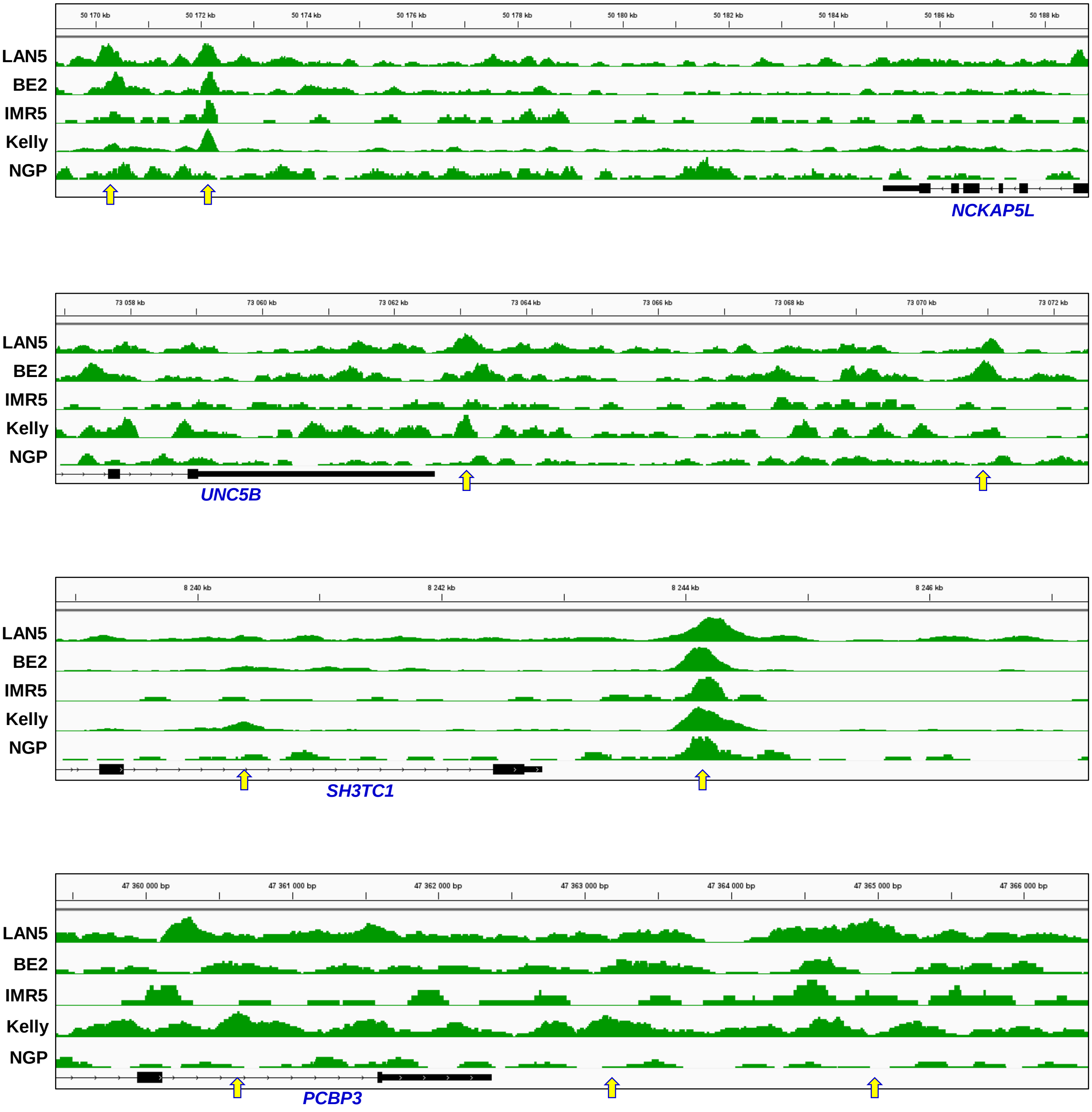
